# Hypoxia and TNF-alpha modulate extracellular vesicle release from human induced pluripotent stem cell-derived cardiomyocytes

**DOI:** 10.1101/2024.02.01.578434

**Authors:** Margarida Viola, Maarten P. Bebelman, Renee G. C. Maas, Frederik J. Verweij, Cor S. Seinen, Saskia C. A. de Jager, Pieter Vader, D. Michiel Pegtel, Joost P. G. Sluijter

## Abstract

Extracellular vesicles (EVs) have emerged as important mediators of intercellular communication in the heart under homeostatic and pathological conditions, such as myocardial infarction (MI). However, the basic mechanisms driving cardiomyocyte-derived EV (CM-EV) production following stress are poorly understood. In this study, we generated human induced pluripotent stem cell-derived cardiomyocytes (hiPSC-CMs) that express NanoLuc-tetraspanin reporters. These modified hiPSC-CMs allow for robust quantification of CM-EV secretion from small numbers of cells without the need for time-consuming EV isolation techniques. We subjected these cells to a panel of small molecules to study their effect on CM-EV biogenesis and secretion under basal and stress-associated conditions. We observed that EV biogenesis is context-dependent in hiPSC-CMs. Nutrient starvation decreases CM-EV secretion while hypoxia increases the production of CM-EVs in a nSmase2-dependent manner. Moreover, the inflammatory cytokine TNF-α increased CM-EV secretion through a process involving NLRP3 inflammasome activation and mTOR signaling. Here, we detailed for the first time the regulatory mechanisms of EV biogenesis in hiPSC-CMs upon MI-associated stressors.

## Introduction

Myocardial infarction (MI) is the leading cause of death worldwide.^1^ Despite current treatment options^2^, heart failure incidence post-MI is expected to increase in the coming years.^3^ It is evident that there is an unmet need for new approaches to promote cardiac repair. Extracellular vesicles (EVs) released by the ischemic heart tissue have recently been reported to drive cardiac repair following acute MI and chronic ischemic remodeling.^4–9^ EVs are membrane particles that mediate intercellular communication by transporting biomolecules, such as proteins and small non-coding RNA, between cells.^10^ Cardiomyocyte-derived EVs (CM-EVs) influence cardiac repair by controlling fibrosis^4^, promoting angiogenesis^5,11^, stimulating macrophages^6^, and regulating endocrine dysfunction in adipocytes^7^. After MI, circulating EVs mobilize bone marrow progenitor cells^9^ and modulate the inflammatory response.^8^ Finally, CM-EVs released by human induced pluripotent stem cell-derived cardiomyocytes (hiPSC-CMs) have therapeutic potential for treating MI. Preclinical studies in rodents showed that these EVs improved heart function when injected directly into infarcted hearts^12–14^ or delivered in a hydrogel.^15,16^

EVs are a heterogeneous group of vesicles of different sizes, ranging from 50 nm to several μm in size, which are produced through various biogenesis mechanisms. They can originate intracellularly from the fusion of multivesicular bodies (MVBs) with the plasma membrane, called exosomes, or be generated by outward budding of the plasma membrane as ectosomes.^17^ The regulation of EV secretion is highly cell-type and context-specific, with microenvironmental cues having strong influences on EV type, quantity, and cargo.^18^ MI has been reported to increase cardiac EV^8,19^ release and change EV cargo, such as microRNA content.^4,5,7^ However, the regulatory mechanisms and EV biogenesis pathways that underlie these changes in CM-EV production following MI are poorly understood. This lack of understanding hampers the potential development of therapeutic strategies aimed at modulating CM-EV production to improve cardiac repair following MI and the design of more effective CM-EV therapies.^20^

The current gap in our understanding of CM-EV secretion is a consequence of the technical difficulties associated with mechanistic EV studies.^17^ The initial step of most EV research typically involves the isolation of EVs using techniques such as differential ultracentrifugation (dUC) and size exclusion chromatography (SEC), but these require a large starting volume of conditioned media and number of cells, are time-consuming, and can lead to co-isolation of protein aggregates.^21,22^ Several strategies have been developed in the last years to study EV production from small cell numbers without the need for EV isolation. Reporters based on tetraspanins and other EV cargo proteins fused to the bright luciferases NanoLuc^23^ and ThermoLuc^24^ have been used to study the mechanisms underlying EV biogenesis directly^25–27^, to identify novel EV modulator drugs^28^, and to optimize culture conditions for the production of therapeutic EVs.^29^ Importantly, most CM-EV secretion studies are currently based on immortalized cell lines^5,6,30–34^ and primary culture^4,7,30,35–38^ of CMs of rodent origin, which are more available, but do not reflect human cardiomyocyte biology. hiPSC-CMs offer this and several other advantages^39^, including their compatibility with genetic interventions for more mechanistic studies.

In this study, we aimed to understand the regulatory mechanisms involved in CM-EV biogenesis and secretion under basal culture conditions and in combination with known post-MI stressors, such as nutrient and oxygen deprivation, and exposure to pro-inflammatory stimuli. To achieve this, we generated hiPSC-CMs expressing NanoLuc-tetraspanin reporters, which enabled the study of CM-EV biogenesis directly from the conditioned media. We identified several pathways involved in CM-EV biogenesis during basal culture conditions using small molecule inhibitors. Furthermore, we found that conditions associated with MI modulate EV release and discovered a role for neutral sphingomyelinase 2 (nSMase2) in hypoxia-driven EV secretion and a potential role for the NLRP3 inflammasome and mTOR signalling in TNF-α-induced EV release.

## Results

### hiPSC-derived cardiomyocytes (hiPSC-CMs) secrete small extracellular vesicles (sEVs)

hiPSC-CMs are increasingly recognized as a relevant *in vitro* model to study human cardiomyocyte biology. In addition, these cells release EVs^12,13,15,19,40–46^, although the variations in the heterogeneity of these EVs remain unclear. The protocol applied here^47^ enabled the generation of a pure population of beating cardiomyocytes (Figure S1, Video S1). To subsequently investigate if the hiPSC-CMs generated secrete small EVs (sEVs), we isolated EVs from the conditioned media using dUC and analyzed the EV pellet following 70 min centrifugation at 100Kx*g* (100K pellet), according to MISEV2018 guidelines.^48^ Western blot confirmed the enrichment relative to cell lysates of classical sEV markers, such as the tetraspanins CD9, CD63, and CD81^49^, syntenin^50^, as well as the presence of ectosome marker Annexin A1.^51^ Depletion of GAPDH, the endoplasmic reticulum marker Calnexin, and the Golgi marker GM130 demonstrated the purity of the isolated sEVs (Figure 1A). Nanoparticle tracking analysis (NTA) (Figure 1B) and transmission electron microscopy (Figure 1C) of the 100K pellet revealed the presence of membrane vesicles with a diameter ranging from 50 to 200 nm and a mean/mode of approximately 90 nm, as expected for sEVs.^52^ Overall, these data confirm that the hiPSC-CMs being used in this research release sEVs.

**Figure 1.**
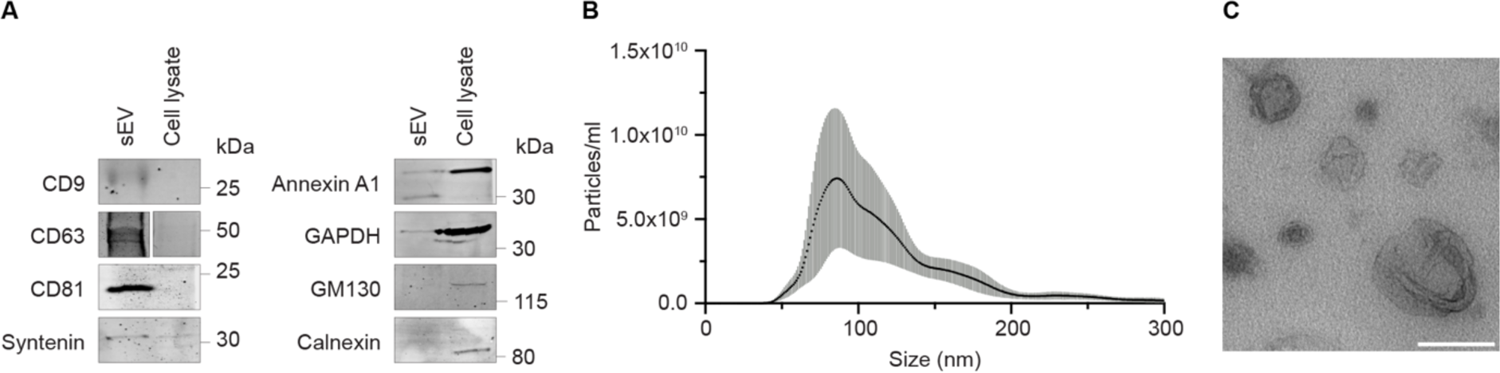
hiPSC-derived cardiomyocytes (hiPSC-CMs) secrete small extracellular vesicles (sEVs). **A.** Western blot analysis of sEVs isolated by differential ultracentrifugation from the conditioned media of hiPSC-CMs for common EV markers (CD9, CD63, CD81, Syntenin and Annexin A1) and cell-enriched markers (GAPDH, GM130 and Calnexin) **B.** Size distribution profile of sEVs analyzed by nanoparticle tracking analysis (NTA). Results are plotted as mean ± SEM of three independent sEV isolations. **C.** Representative transmission electron microscopy (TEM) image of sEVs (scale bar = 200 nm).

### NanoLuc-tetraspanin reporters enable direct quantification of sEV secretion from hiPSC-CMs

To study CM-EV secretion directly without the need for EV isolation, we generated hiPSC-CMs stably expressing the tetraspanins CD9, CD63, and CD81 tagged at their N-termini with NanoLuc and a hemagglutinin (HA)-tag to facilitate antibody-based detection (Figure 2A). Transduction efficiency and location of the HA-NanoLuc-(NL)-tetraspanin reporters were verified by immunofluorescence microscopy (Figure 2B). HA-NL-CD9 and HA-NL-CD81 were localized to the plasma membrane and endosomal compartments, whereas HA-NL-CD63 was mostly present in endosomal compartments. This is consistent with their endogenous localization (Figure S2). Upon addition of the NanoLuc substrate Furimazine, we could reliably measure luminescence in both cells and conditioned culture supernatant of hiPSC-CMs (Figure 2C). To confirm that the NanoLuc activity in the culture supernatant originated from EV-associated HA-NL-tetraspanin reporters, we separated EVs and soluble proteins using SEC and measured NanoLuc activity in each fraction (Figure 2D). NanoLuc activity peaked in EV-enriched fractions 8-14, which contain the most particles as confirmed by NTA (Figure 2E), and was nearly absent in fractions 20-26, which are enriched in soluble proteins and extracellular particles that are smaller than the 70 nm pore size of the SEC column (Figure 2F).

**Figure 2.**
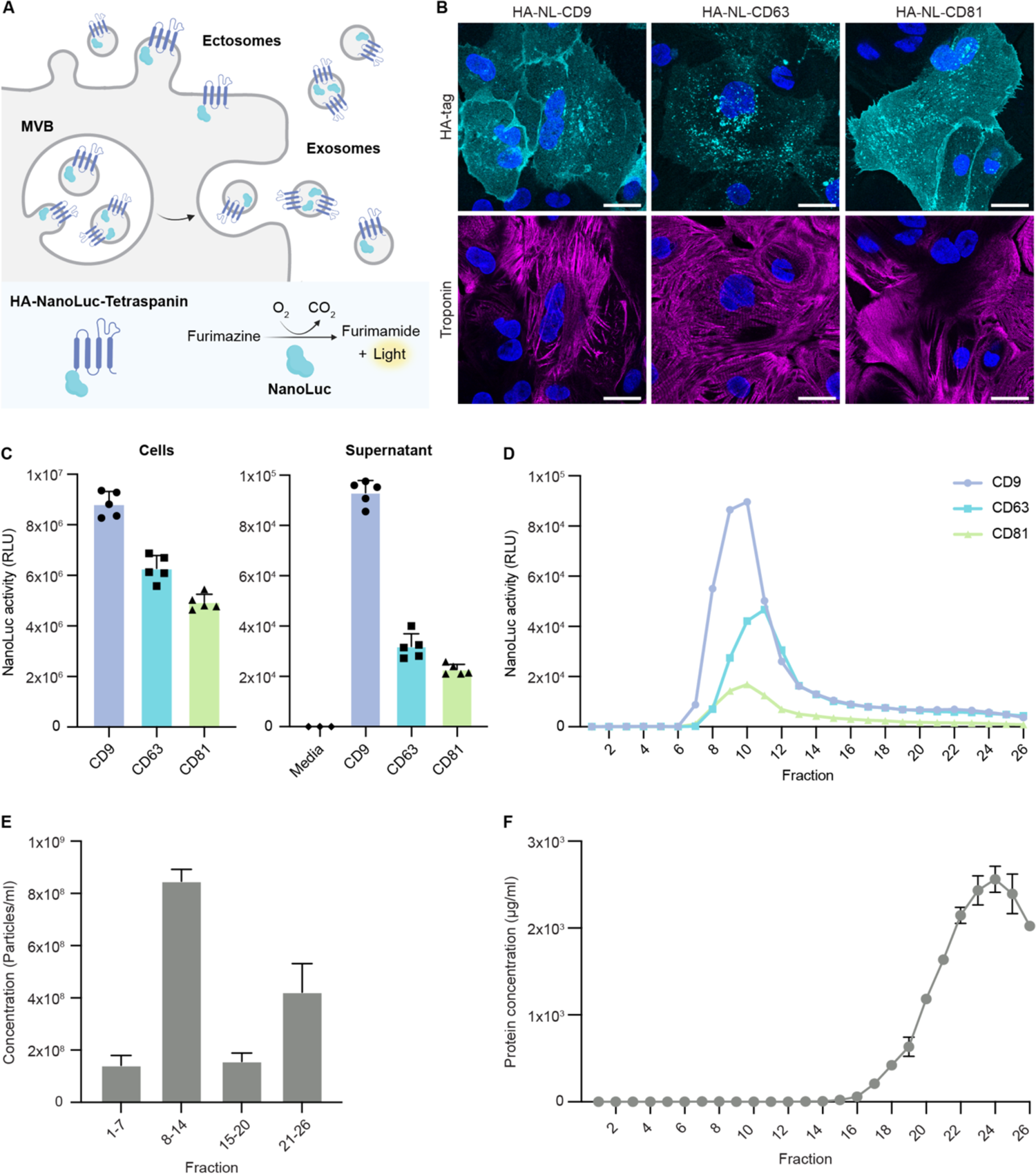
NanoLuc-tetraspanin reporters enable direct quantification of sEV secretion from hiPSC-CMs. **A.** Schematic representation of sEV quantification using HA-NanoLuc(NL)-tetraspanin reporters. HA-NL-Tetraspanin reporters are sorted into exosomes and ectosomes and tetraspanin-positive EV secretion can be quantified by measuring NanoLuc bioluminescent activity in the culture supernatant. **B.** Representative immunofluorescence images of hiPSC-CMs transduced with HA-tagged NL-tetraspanin reporters and stained for HA-tag and cardiac troponin (scale bar = 20 µm). **C.** Representative measurement of NanoLuc activity in cells and conditioned culture supernatant of hiPSC-CMs expressing HA-NL-tetraspanin reporters cultured in a 96-well format. Data is represented as mean ± SD. **D.** Representative luminescence measurement of size exclusion chromatography (SEC) fractions obtained from the supernatant of hiPSC-CMs expressing HA-NL-tetraspanin reporters. **E.** Representative graph of particle number (> 50 nm) of pooled SEC fractions analyzed by nanoparticle tracking analysis (NTA) with mean ± SEM. **F.** Representative graph of protein concentration of SEC fractions with mean ± SD.

In summary, we generated hiPSC-CMs that stably express HA-NL-tetraspanin reporters. These cells can be utilized to measure tetraspanin-positive EV release directly from the conditioned medium of a relatively small number of cells.

### Screening of chemical modulators identifies pathways involved in extracellular vesicle biogenesis of hiPSC-CMs

To improve our understanding of CM-EV biogenesis mechanisms, we subsequently used hiPSC-CMs expressing HA-NL-tetraspanin reporters to screen a range of small molecules known to modulate EV secretion in other cell types (Table S1).^26,53^ The screen was performed in 96-well format and the NanoLuc activity in the collected culture supernatant was normalized to the corresponding cellular NanoLuc activity, to account for small differences in the plating of the cells (Figure 3A). Small molecule concentrations were selected based on the literature and alamarBlue assay was used to ensure that the cell viability was above 80% (Figure S3). As a positive control, we included the vacuolar proton pump H+ ATPase (vATPase) inhibitor Bafilomycin, which has been reported to strongly increase sEV secretion in multiple cell types.^25,26,54^ Indeed, we find that Bafilomycin increased the secretion of all three HA-NL-tetraspanins from hiPSC-CMs (Figure 3B). Similarly, the actin polymerization inhibitor cytochalasin D, the calpain inhibitor calpeptin, and the PIKfyve inhibitor apilimod increased HA-NL-tetraspanin secretion (Figure 3C-E). In contrast, the ROCK inhibitor Y-27632 significantly reduced HA-NL-CD81 secretion by ±29% and showed a similar trend for CD9 and CD63. The neutral sphingomyelinase 2 (nSMase2) inhibitor GW4869, the Niemann-Pick disease type C1 (NPC1) inhibitor U18666A, the Rab27 inhibitor nexinhib20, the VPS34 inhibitor SAR405, the PI3K inhibitor wortmannin and the calcium ionophore A23187 did not significantly affect HA-NL-tetraspanin secretion by cardiomyocytes. These findings confirm that cardiomyocytes expressing HA-NL-tetraspanins can be used to screen for modulators of CM-EV secretion and suggest that multiple players participate in the biogenesis of CM-EVs.

**Figure 3.**
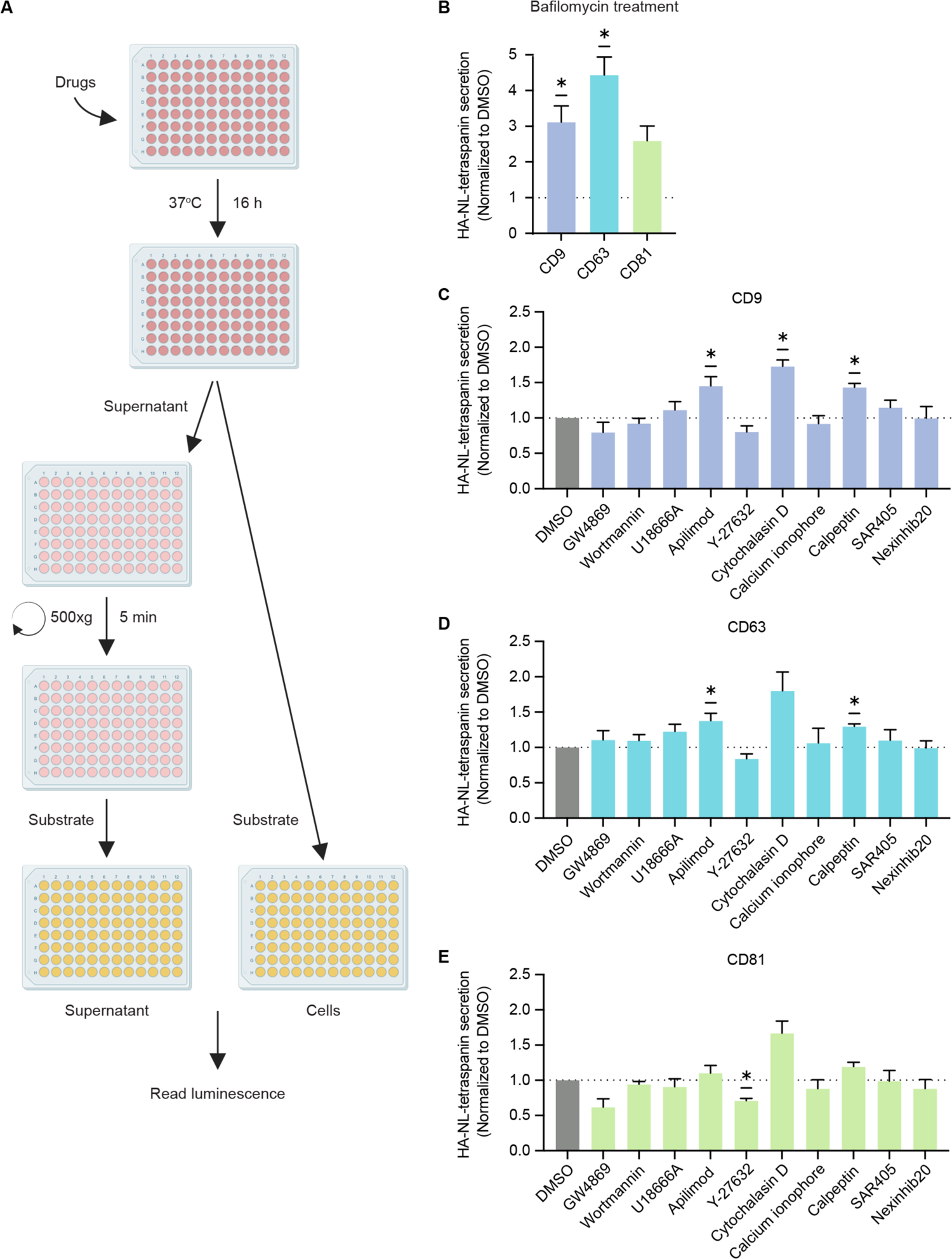
Effect of extracellular vesicle biogenesis modulators on NanoLuc-tetraspanin secretion by hiPSC-CMs. **A.** Experimental method schematic of a small molecule screen using NanoLuc-tetraspanin reporters in hiPSC-CMs. **B.** NanoLuc-tetraspanin secretion upon bafilomycin treatment normalized to the DMSO control. **C-E.** NanoLuc-tetraspanin secretion of CD9 (B), CD63 (C) and CD81 (D) analysis using various small molecules involved in sEV secretion: GW4869 (5 μM), wortmannin (0.1 μM), U18666A (5 μM), apilimod (0.5 µM), Y-27632 (5 μM), cytochalasin D (0.1 μM), calcium ionophore A23187 (1 μM), calpeptin (5 µM), SAR405 (1 μM), nexinhib20 (1 μM) and bafilomycin (0.1 µM). Results from B to E are plotted as mean ± SEM of three independent experiments. Statistical analysis was done using a one-sample, two-tailed T-test against a theoretical mean of 1. *p <0.05.

### Stress conditions modulate extracellular vesicle secretion by hiPSC-CMs

MI is characterized by physiological stress due to a lack of blood supply, limiting nutrient and oxygen levels (hypoxia) in cardiac tissue. When ischemic cardiac cells start to die, they release inflammatory cytokines such as tumor necrosis factor-alpha (TNF-α), initiating inflammation.^55^ In addition, levels of the vasoconstrictor hormone angiotensin II (Ang II) are increased in response to decreased blood pressure during MI.^56^ Hypoxia, nutrient starvation, TNF-α, and Ang II have all been linked to changes in EV biogenesis and cargo selection in cardiomyocytes^19,30,57^ or other cell types^58–61^.

To gain insight into the underlying mechanisms, we first exposed hiPSC-CMs expressing HA-NL-tetraspanins to cardiac stressors and assessed the effect of these on HA-NL-tetraspanin release after 24 and 48 hours (Figure 4A-C). Nutrient starvation reduced the secretion of all HA-NL-tetraspanin reporters already after 24h, whereas hypoxic culture conditions strongly increased the secretion of HA-NL-CD63 and HA-NL-CD81 after 48 hours, but not HA-NL-CD9 (Figure 4A). Interestingly, TNF-α treatment resulted in a time-dependent increase in HA-NL-tetraspanin secretion. Importantly, at the concentration used in this study, TNF-α does not induce CM cell death^62^, as was confirmed using a cell viability assay (Figure S3). Cardiomyocytes are known to express TNF-α receptors I and II^63^, and neutralizing antibodies against both receptors significantly reduced NanoLuc-CD81 secretion during the presence of TNF-α (Figure 4D), suggesting that both TNF receptors contribute to TNF-α-induced CD81-positive EV secretion.

**Figure 4.**
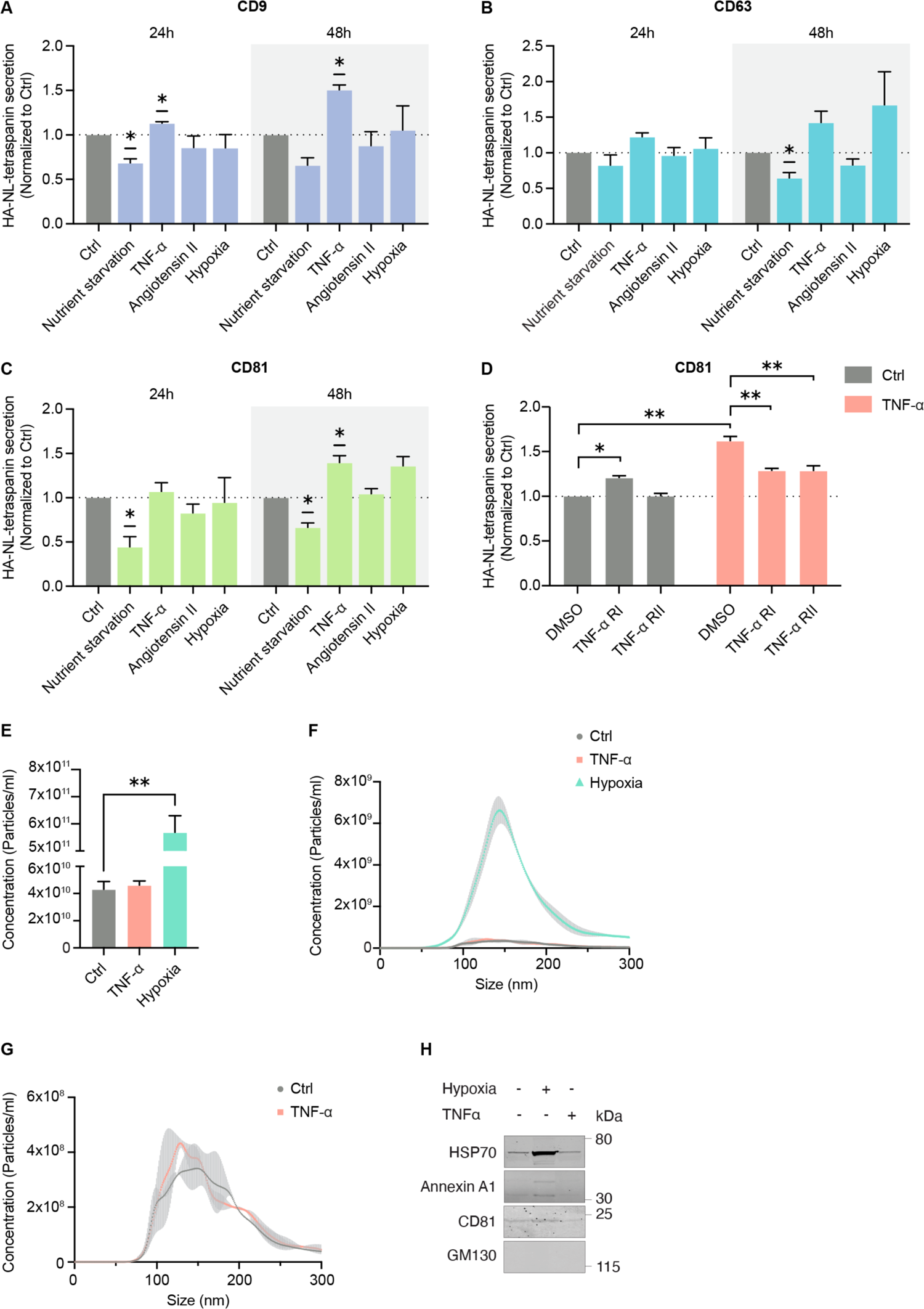
Stress conditions modulate extracellular vesicle secretion by hiPSC-CMs. **A-C.** Effect of various stress conditions (nutrient starvation, 10 ng/ml TNF-α, 1 µM angiotensin II, and hypoxia) on the secretion of NanoLuc-CD9 (A), NanoLuc-CD63 (B) and NanoLuc-CD81 (C) after 24 and 48 hours. Results are plotted as mean ± SEM of three independent experiments. Statistical analysis was done using a one-sample, two-tailed T-test against a theoretical mean of 1. *p <0.05. **D.** NanoLuc-CD81 secretion under control (Ctrl) and TNF-α conditions in combination with neutralizing antibodies against TNF-α receptor I (TNF-α RI) or II (TNF-α RII). Results are plotted as mean ± SEM of three independent experiments. Comparisons in the control group were analyzed using a one-sample, two-tailed T-test against a theoretical mean of 1. Comparisons within the TNF-α group were analyzed using a one-way ANOVA with Dunnett correction against the DMSO condition. *p <0.05, **p <0.01. **E-G.** Particle number (E) and size distribution profiles (F, G) of sEVs released after 48 hours under control, TNF-α, and hypoxia were analyzed by nanoparticle tracking analysis (NTA). Results are plotted as mean ± SEM of two independent experiments and data was analyzed using a one-way ANOVA with Dunnett correction against the control condition. **p <0.01. **H.** Representative western blot analysis of two independent experiments of sEVs released after 48 hours under control, hypoxia, and TNF-α for EV markers (HSP70, annexin A1, and CD81) and the cell-enriched marker GM130. sEV samples from equal cell numbers and conditioned medium volumes were loaded.

To validate the effects of TNF-α and hypoxia on total CM-EV secretion we isolated CM-EVs by dUC from the conditioned medium of hiPSC-CMs cultured under hypoxic conditions or treated with TNF-α for 48h and measured the particle number and size distribution using NTA (Figure 4E-G). We found that hypoxia significantly increased the particle number compared to control conditions, whereas there was no effect of TNF-α on bulk EV secretion. The discrepancy of results between NTA and the HA-NL-tetraspanin secretion assay suggests that, in addition to stimulating the secretion of tetraspanin-positive EVs, hypoxia also stimulates the release of tetraspanin-negative EVs and/or non-vesicular extracellular particles. In contrast, TNF-α seems to induce the release of tetraspanin-positive EVs, but not to an extent that is detectable in the bulk EV population. This shows that NL-tetraspanin reporters are very sensitive to small changes in EV release. Western blot analysis of EVs isolated from equal cell numbers and culture medium volume confirmed increased levels of the EV markers HSP70, annexin A1, and CD81 in the sEV pellet of hypoxic cells (Figure 4H). Taken together, these findings highlight that EV biogenesis is highly context-dependent in hiPSC-CMs. Hypoxia induces a significant increase in the overall EV release, whereas TNF-α specifically promotes the secretion of tetraspanin-positive EVs.

### Neutral sphingomyelinase 2 mediates extracellular vesicle release under hypoxic conditions

To further explore the potential mechanisms underlying TNF-α- and hypoxia-stimulated EV production, we investigated the effect of EV biogenesis-modulating small molecules on HA-NL-tetraspanin release (Figure 5A-C). Under basal culture conditions, the effects of these modulators were similar between 16 and 48 hours (Figure 3B-D), except for the NPC1-inhibitor U18666A, which significantly increased HA-NL-CD9 and -CD81 secretion, and for calpeptin, which did not affect HA-NL-tetraspanin secretion after 48 hours.

**Figure 5.**
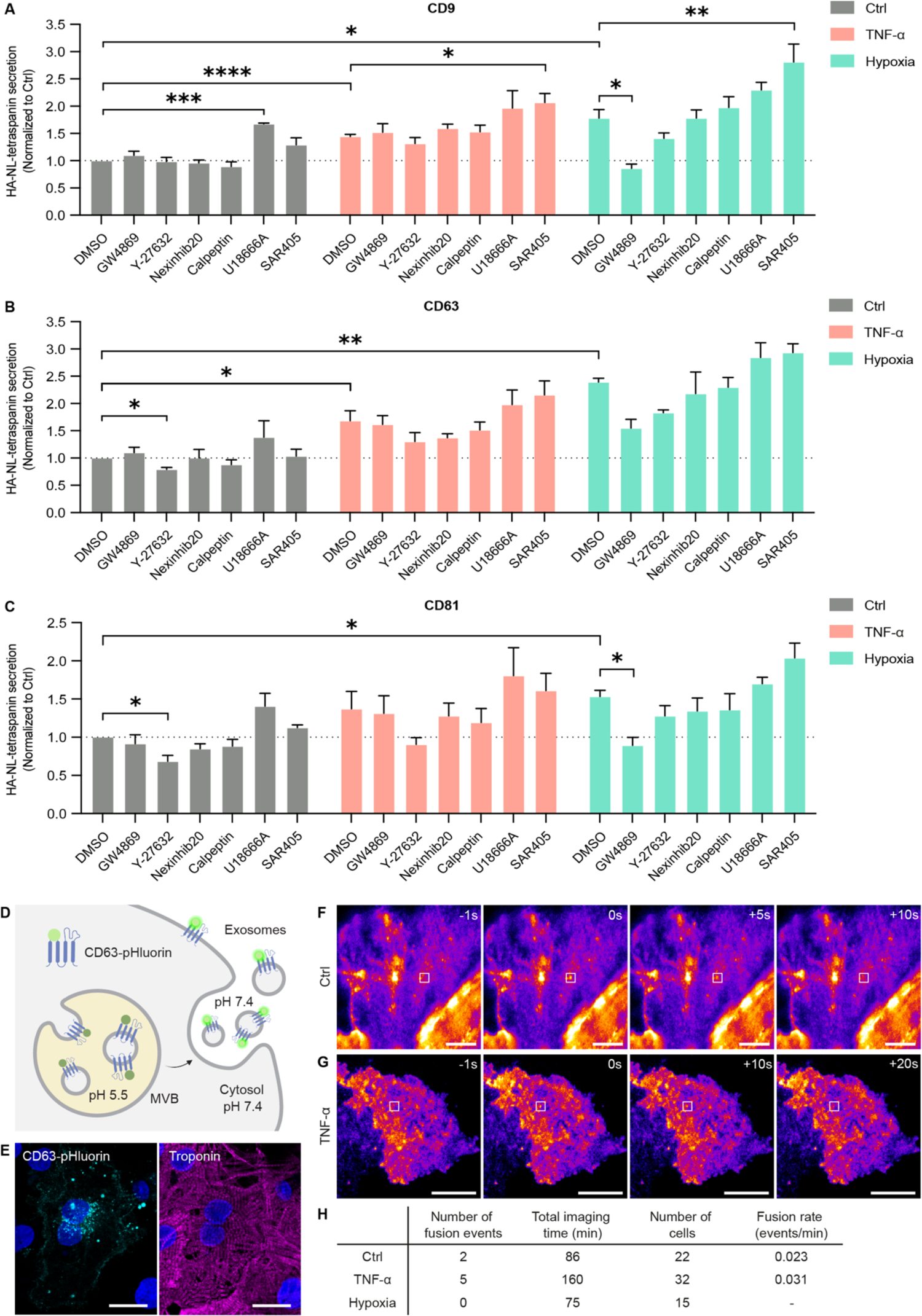
nSmase2 mediates extracellular vesicle release under hypoxic conditions. **A-C.** Effect of EV biogenesis modulators GW4869 (5 μM), Y-27632 (5 μM), nexinhib20 (1 μM), calpeptin (5 µM), U18666A (5 μM), and SAR405 (1 μM) on NanoLuc-tetraspanin secretion under control (Ctrl), 10 ng/ml TNF-α and hypoxic conditions for CD9 (A), CD63 (B), CD81 (C) positive EVs after incubation for 48 hours. Results are plotted as mean ± SEM of three independent experiments. Comparisons to control conditions were analyzed using a one-sample, two-tailed T-test against a theoretical mean of 1. Comparisons within the TNF-α and hypoxia groups were analyzed using a one-way ANOVA with Dunnett correction against the DMSO condition within each group. *p <0.05, **p <0.01. **D.** Schematic representation of CD63-pHluorin for imaging of MVB-plasma membrane fusion. **E.** Localization of CD63-pHluorin in hiPSC-CMs (scale bar = 20 µm). **F, G.** Time-lapse TIRF imaging of a CD63-pHluorin fusion event in control (F) and TNF-α-treated (G) hiPSC-CMs. **H.** MVB-plasma membrane fusion rate in CD63-pHluorin hiPSC-CMs under control, TNF-α, and hypoxic conditions of two independent experiments for each condition.

The ROCK inhibitor Y-27632 significantly decreased HA-NL-CD63 and -CD81 secretion in control conditions and showed a similar trend for all HA-NL-tetraspanins upon TNF-α treatment or hypoxic conditions. Interestingly, whereas nSmase2 inhibition by GW4869 did not decrease EV secretion in control conditions or upon TNF-α treatment, it significantly inhibited HA-NL-CD9 and -CD81 secretion under hypoxic conditions. A similar trend was observed for HA-NL-CD63 secretion. Thus, it appears that the mechanisms underlying EV biogenesis in hiPSC-CMs are context-dependent.

The contribution of nSmase2 to HA-NL-tetraspanin secretion under hypoxic conditions suggests that hypoxia drives the release of MVB-derived exosomes.^64,65^ This speculation agrees with previous studies revealing that hypoxia induces peripheral trafficking of late endosomal compartments, a limiting step in exosome secretion.^66,67^ To investigate whether the TNF-α- and hypoxia increase exosome secretion specifically, we first assessed the effect of these treatments on MVB localization using IF microscopy against the MVB marker CD63. We did not observe differences in MVB localization when compared to control hiPSC-CMs (Figure S4). Due to their overlapping content and biophysical properties, most EV isolation and characterization strategies, including HA-NL-tetraspanin-based approaches, do not formally distinguish between exosomes and ectosomes. To gain a more conclusive answer, we transduced cardiomyocytes with a CD63-pHluorin reporter construct, which enables quantification of MVB exocytosis rate as a proxy for exosome release (Figure 5D).^65,68^ Fluorescence microscopy confirmed the presence of CD63-pHluorin in intracellular compartments, in line with its incorporation into MVBs (Figure 5E). Live cell total internal reflection fluorescence (live-TIRF) microscopy revealed several sudden bursts in fluorescence intensity corresponding to the fusion of CD63-pHluorin-containing MVBs with the plasma membrane (Figure 5F-G, Video S2-3).^68^ Possibly because the fusion rate under control conditions and upon TNF-α treatment is very low in hiPSC-CMs, we failed to measure a significant increase in MVB exocytosis upon TNF-α exposure (Figure 5H). Importantly, we observed no fusion events under hypoxic conditions, suggesting that hypoxia does not lead to a large increase in MVB exocytosis rate and exosome release.

### Inflammasome activation and mTOR signaling contribute to TNF-alpha-induced HA-NL-CD81 secretion

TNF-α signaling upregulates the expression of inflammasome components and hypoxia-induced mitochondrial dysfunction can activate the inflammasome, leading to the production of cytokines, such as interleukin-1 beta (IL-1β) and interleukin-18 (IL-18).^69,70^ Several studies have linked inflammasome activation to elevated EV secretion.^71–73^ To investigate the role of the inflammasome in TNF-α- and hypoxia-induced EV secretion, we inhibited the inflammasome sensor NACHT, leucine-rich repeat, and pyrin domain–containing protein 3 (NLRP3) using the small molecule MCC950.^74^ TNF-α-induced, but not basal or hypoxia-induced, HA-NL-CD81 secretion was significantly inhibited by MCC950 and a similar trend was observed for HA-NL-CD63 (Fig 6A-B). Considering that TNF-α signaling can trigger the mechanistic target of rapamycin (mTOR) activation^75^, and given the role of mTOR signaling in inflammasome formation^76^, we also tested the effect of the mTOR inhibitor rapamycin and found that it inhibited TNF-α-induced HA-NL-CD81 secretion (Fig 6A-B). In conclusion, our results suggest a role for inflammasome activation and mTOR signaling in TNF-α-induced CM-EV secretion.

**Figure 6.**
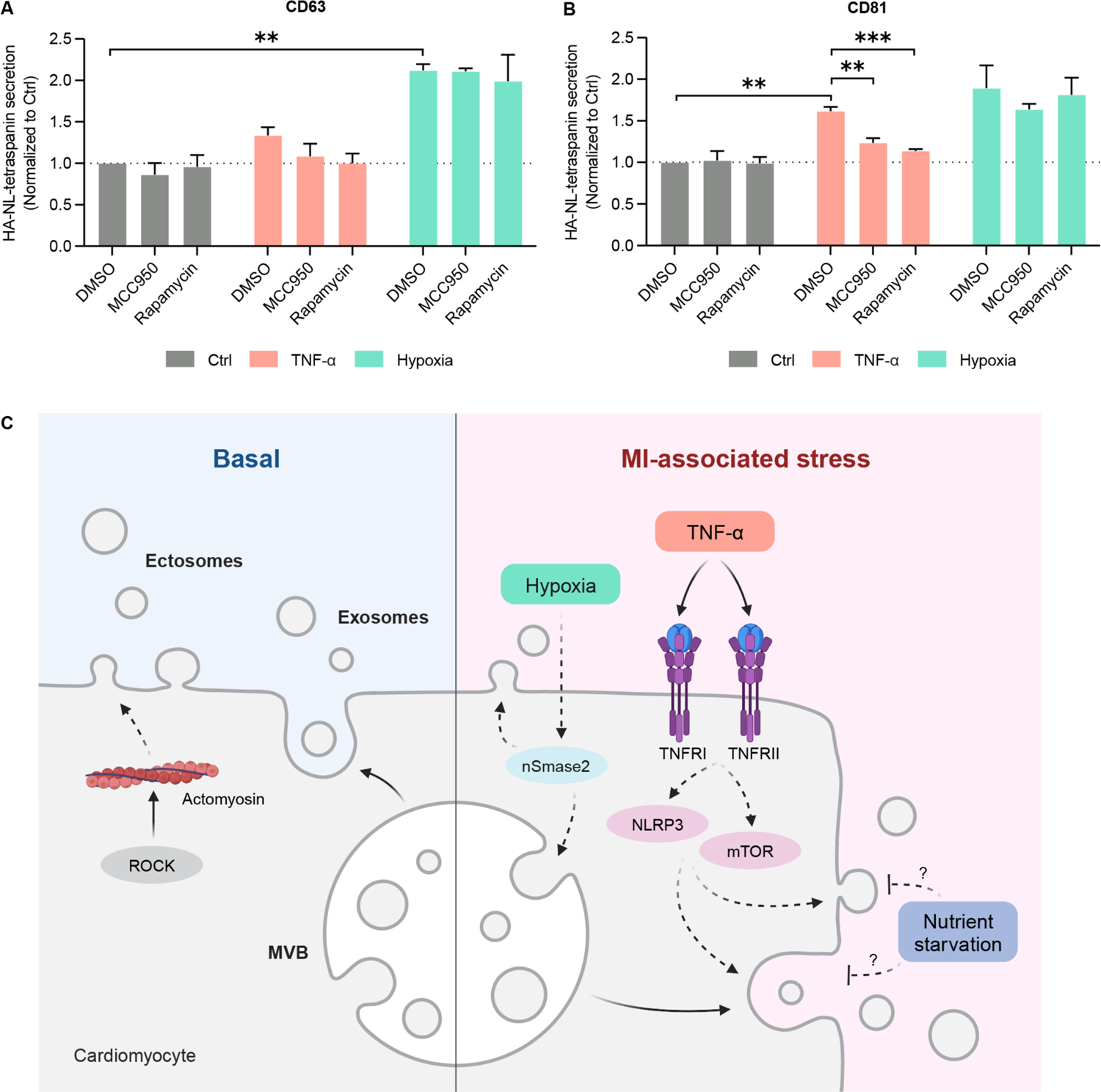
Inflammasome activation and mTOR signaling contribute to TNF-alpha-induced HA-NL-CD81 secretion. **A, B.** NanoLuc-tetraspanin secretion of CD63 (A) and CD81 (B) after incubation with the inflammasome inhibitor MCC950 or mTOR inhibitor rapamycin under control, TNF-α and hypoxic conditions. Results are plotted as mean ± SEM of three independent experiments. Comparisons to control conditions were analyzed using a one-sample, two-tailed T-test against a theoretical mean of 1. Comparisons within the TNF-α and hypoxia groups were analyzed using a one-way ANOVA with Dunnett correction against the DMSO condition within each group. *p <0.05, **p <0.01. **C.** Scheme summarizing the findings of this study. EV biogenesis is context-dependent in hiPSC-CMs. Under basal culture conditions, hiPSC-CMs can secrete exosomes, and ectosomes, which release is mediated in part by ROCK. Nutrient starvation decreases CM-EV secretion while hypoxia increases the production of CM-EVs in a nSmase2-dependent manner. TNF-α increased CM-EV secretion through a process involving NLRP3 inflammasome activation and mTOR signaling. Abbreviations: MI, myocardial infarction; ROCK, Rho-associated protein kinase; MVB, multivesicular body; nSMase2, neutral sphingomyelinase 2; TNF-α, tumor necrosis factor alpha; TNFRI, TNF-α receptor 1; TNFRII, TNF-α receptor 2; NLRP3, NLR family pyrin domain containing 3; mTOR, mammalian target of rapamycin.

## Discussion

EVs have emerged as important mediators of intercellular communication in the heart under homeostatic and pathological conditions, such as MI. However, the biogenesis mechanisms that contribute to CM-EV secretion under these conditions are poorly understood. This hampers the discovery of new therapeutic targets to modulate CM-EV production and the design of more effective CM-EV therapies to promote cardiac repair after MI. Here, we generated HA-NL-tetraspanin-expressing hiPSC-CMs that enabled us to robustly quantify EV secretion from small numbers of cells. Using this approach, we identified modulators of CM-EV biogenesis under basal and stress-associated conditions (Figure 6C).

To study EV biogenesis in hiPSC-CMs we used a panel of small molecules that target known EV biogenesis pathways (Table S1) and observed that the ROCK inhibitor Y-27632 decreased CM-EV secretion. ROCK has been reported to participate in ectosome release by inducing outward budding of the plasma membrane via actomyosin remodeling.^77,78^ There was no significant effect on CM-EV secretion upon the inhibition of nSmase2, which contributes to ceramide-dependent intraluminal vesicle budding and MVB formation.^64^ Similarly, we did not observe any effect of inhibitors of the endosomal cholesterol-transporter NPC1 or the small GTPase Rab27, which contribute to exosome secretion by controlling MVB positioning and docking to the plasma membrane, respectively.^79,80^ These findings suggest that, under basal culture conditions, hiPSC-CMs release more ectosomes than exosomes. In line with this, live TIRF imaging with CD63-pHluorin revealed that hiPSC-CMs have a very low MVB-plasma membrane fusion rate compared to other cell types.^68^ hiPSC-CMs are not unique in their preference for ectosome secretion. Recent studies have reported that the majority of sEVs secreted by HEK293 and HeLa cells originate from the plasma membrane.^25,81,82^ Similar to our findings in hiPSC-CMs, NanoLuc-CD63 secretion from HEK293 cells was not inhibited by GW4869 or U1866A treatment^25,26^, except when bafilomycin was used to induce exosome secretion.^25^

Additionally, we tested other small molecules that have been reported to modulate EV production. mTOR signaling and autophagy can both inhibit and stimulate the release of sEVs, depending on cell type and cellular context.^83–85^ Our results suggest that mTOR signaling and canonical autophagy do not play a role in tetraspanin-positive EV release from hiPSC-CMs under basal conditions, as shown using rapamycin and SAR405. However, we found that the PIKfyve inhibitor apilimod, a stimulator of secretory autophagy and exosome secretion in a prostate cancer cell line^86^, increased CM-EV release. The actin polymerization inhibitor cytochalasin D also increased HA-NL-tetraspanin secretion from hiPSC-CMs, consistent with previous observations in HEK293 cells.^26^ Finally, we found that 16h treatment with the calpain inhibitor calpeptin induced HA-NL -tetraspanin release, whereas it is described to decrease EV secretion by other cell types.^87,88^. Interestingly, this stimulatory effect was lost after 48h, suggesting that short-term calpeptin treatment leads to a temporary surge in CM-EV secretion.

The rate of EV secretion and the EV biogenesis mechanisms highly depend on the cellular context.^18^ We sought to explore the effect of MI-associated stress factors on human CM-EV secretion. Nutrient starvation decreased HA-NL-tetraspanin secretion by hiPSC-CMs. This is consistent with observations in other cell types^25,58^, but different from a previous finding in immortalized H9C2 embryonic cardiomyocytes, where glucose starvation increased EV secretion.^31^ The decrease in EV secretion upon nutrient starvation may be due to the induction of autophagy and redirection of MVBs into the autophagic pathway, resulting in decreased EV secretion.^59^ MI increases cardiac levels of Ang II, a hormone that has been described to increase EV production by HEK293T cells and neonatal and adult rat cardiomyocytes.^57,61,89^ However, we did not find differences in HA-NL-tetraspanin secretion upon stimulation of hiPSC-CMs with Ang II. Consistent with previous work, we observed that hypoxia increased the production of tetraspanin-positive CM-EVs.^19,42,57,90^ Notably, several other studies did not find a significant effect of hypoxia on CM-EV secretion. These discrepancies may be attributed to variations in the origin of the cardiomyocytes, EV isolation and quantification methodology, and the duration and method used to induce hypoxia.^30,45,90^ The cytokine TNF-α, which contributes to the inflammatory response following MI, has been reported to stimulate EV release from tumor cells and astrocytes^60,91^, but its role in CM-EV secretion remained unknown. Our results suggest that stimulation with a non-toxic concentration of TNF-α increases sEV secretion by cardiomyocytes.

To shed light on the mechanisms underlying hypoxia- and TNF-α-induced EV secretion, we tested the panel of EV-modulating small molecules on HA-NL-tetraspanin secretion under these stress conditions. We found that the nSmase2 inhibitor GW4869 blocked hypoxia-induced EV secretion, indicating that hypoxia might increase exosome secretion. However, we did not observe any MVB-plasma membrane fusion events using CD63-pHuorin following hypoxia treatment. The CD63-pHluorin imaging data needs to be interpreted with care since the low fusion activity in hiPSC-CMs precluded an accurate assessment of differences in MVB exocytosis rate. Nevertheless, the complete absence of fusion events, and the lack of Rab27 and NPC1-involvement in hypoxia-induced HA-NL-tetraspanin secretion, argue against a strong induction of exosome secretion under hypoxic conditions. Further research is needed to understand the membrane origin of hypoxia-induced nSmase2-dependent EV secretion by hiPSC-CMs, especially given that nSmase2 has also been demonstrated to regulate ectosome release.^92^ None of the EV-modulating drugs significantly inhibited TNF-α-stimulated secretion, although the ROCK inhibitor Y-27632 appeared to reduce TNF-α-stimulated and basal CM-EV secretion to a similar extent. TNF-α primes the NLRP3 inflammasome and activates mTOR signaling^69,75^, and both signaling pathways have been linked to enhanced EV secretion.^71,84^ Indeed, our results suggest that NLRP3 inflammasome and mTOR signaling contribute to the TNF-α-induced secretion of CD81-positive EVs. Overall, our findings suggest that hypoxia and TNF-α induce CM-EV secretion through different mechanisms and highlight the power of HA-NL-tetraspanin hiPSC-CMs to investigate CM-EV biogenesis under control and MI-associated conditions.

A major advantage of HA-NL-tetraspanin reporters is that they can be used to screen for modulators of EV secretion in a high-throughput manner. For example, HA-NL-tetraspanin reporters have been used to screen a kinase inhibitor library in HEK293 cells^25^ and an FDA-approved drug library in a breast cancer cell line.^28^ Alternatively, small interfering RNA (siRNA) and parallel CRISPR screens may be used to study genes involved in CM-EV generation. HA-NL-tetraspanin reporters can also be used to optimize culture conditions for the production of large numbers of therapeutic hiPSC-CM EVs.^29^ Another exciting possibility is to express HA-NL-tetraspanin reporters in genetically modified or patient-derived hiPSC-CMs that model cardiomyopathies associated with changes in the secretion or function of CM-EVs.^93,94,95^ In this case, HA-NL-tetraspanin reporters could reveal the impact of genetic mutations or specific cardiomyopathy phenotypes on EV generation mechanisms. Finally, EVs isolated from HA-NL-tetraspanin-expressing cardiomyocytes could be used to study CM-EV uptake by other cell types *in vitro*, or even injected into the infarcted mouse heart to study their cardiac retention and biodistribution following MI.^96–98^

In conclusion, we generated HA-NL-tetraspanin-expressing hiPSC-CMs that allowed for robust quantification of CM-EV secretion from small numbers of cells without the need for cumbersome and time-consuming EV isolation techniques. We used these cells to identify modulators of CM-EV biogenesis under basal and MI-associated stress conditions, demonstrating the power of this multiple-reporter approach. We observed that under basal culture conditions, hiPSC-CMs can secrete exosomes, and ectosomes as part of CM-EV release is mediated by ROCK. Additionally, we demonstrated that EV biogenesis is context-dependent in hiPSC-CMs. Nutrient starvation decreased CM-EV secretion while hypoxia increased the production of CM-EVs in a nSmase2-dependent manner. Moreover, TNF-α increased CM-EV secretion through a process involving NLRP3 inflammasome activation and mTOR signaling. This work paves the way for larger screens that will improve our understanding of the molecular mechanisms underlying CM-EV biogenesis in homeostasis and disease, which will aid the development of novel pharmacological strategies for MI.

## Supporting information

Video S1. Morphology of beating hiPSC-CMs 26 days after maturation imaged using a 10x objective.

Video S2. TIRF microscopy of hiPSC-CMs expressing CD63-pHluorin. The video is played at 10x speed. Scale bar 10 um.

Video S3. TIRF microscopy of hiPSC-CMs expressing CD63-pHluorin after 48 hours incubation with TNF-a. The video is played at 10x speed. Scale bar 10um

## Author contributions

MV and MB conceptualization, performance of experiments, data analysis, and writing, RM, FV, and CS performance of experiments, SJ, PV, and MP supervision, JS supervision, and funding acquisition.

## Acknowledgements

All illustrations were designed using BioRender (available at BioRender.com). This work was financially supported by the European Research Council (ERC) Consolidator grant EVICARE (number 725229) to JS. We thank the Cell Microscopy Core of the Department of Cell Biology at the University Medical Center Utrecht, The Netherlands, for granting us access to their microscopes.

## Conflict of Interest

The authors declare no competing interests.

## Materials and methods

### Chemicals

GW4869 (D1692), wortmannin (W3144), U18666A (U3633), apilimod (SML2974), Y-27632 (688001), cytochalasin D (C8273), calcium ionophore A23187 (C7522), nexinhib20 (SML1919), bafilomycin A1 (B1793), MCC950 (538120) and rapamycin (R0395) were purchased from Sigma-Aldrich. Calpeptin (HY-100223) and SAR405 (HY-12481) were obtained from MedChem Express. Recombinant human TNF-α (300-01A) and angiotensin II (A9525) were from PeproTech and Sigma-Aldrich, respectively.

### hiPSC-derived cardiomyocyte differentiation and culture

The hiPSC line was obtained from the Stanford Cardiovascular Institute Biobank (SCVI-273) and non-enzymatically passaged using 0.5 mM EDTA (Thermo Fisher) every 4 days as previously described.^99^ hiPSC cells were maintained in 0.1 mg/mL matrigel-coated 6-well plates (Corning, 356230) in E8 complete medium (Gibco, A1517001) for at least two passages, and grown to ∼90% confluence before starting cardiac lineage differentiations. hiPSCs were efficiently differentiated as previously described^47^, using the RPMI/B27 minus insulin medium (Gibco, A1895601) supplemented with 7–8 µM CHIR99021 (cell line specific) (SelleckChem, S2924) for the first 2 days and 2 µM Wnt-C59 (SelleckChem, S7037) for another 48 hours. The proliferation of hiPSC-CMs can be conducted by removing cell-to-cell contacts and small molecule inhibition of GSK3 with CHIR99021.^100^ Subsequential multiple low-density passaging and reintroduction of 2 µM CHIR99021 to the RPMI/B27 media (Gibco, 17504044) results in an expansion. During the expansion phase, hiPSC-CMs were grown in matrigel-coated T175 flasks and the media was changed every 2-3 days. To mature the CMs, we used a previously described metabolic maturation medium^101^, which consists of DMEM without glucose (Gibco, 11966025) supplemented with 3 mM glucose (Sigma Aldrich, G7021), 10 mM L-lactate (Sigma Aldrich, 71718), 5 μg/ml Vitamin B12 (Sigma Aldrich, V6629), 0,82 μM Biotin (Sigma Aldrich, B4639), 5 mM Creatine monohydrate (Sigma Aldrich, C3630), 2 mM Taurine (Sigma Aldrich, T0625), 2 mM L-carnitine (Sigma Aldrich, C0283), 0,5 mM Ascorbic acid (Sigma Aldrich, A8960), 1x non-essential amino acid solution (Gibco, 11140050), 0.5% (w/v) Albumax (Gibco, 11020021), 1x B27 and 1% KnockOut Serum Replacement (KOSR, Gibco, 10828028). Maturation was carried out for at least 3 weeks with media changes every 3-4 days before proceeding to experiments. Mature hiPSC-CMs were detached by incubation with TrypLE select 10x (Gibco, A1217702) for 15-30 minutes at 37°C. Cells were then re-plated into matrigel-coated 96-well plates to a density of 70 000 cells/well and cultured in RPMI/B27 supplemented with 10% KOSR and 10 μM Y-27632 (Selleck Chemicals, S1049). Media was switched to RPMI/B27 the following day and changed to maturation media 2-3 days later. hiPSC-CMs were kept in maturation media for another 7 days to allow cell recovery before drug testing. All HA-NL-tetraspanin assays were performed in maturation media which was refreshed every 3-4 days.

### EV isolation using differential ultracentrifugation (dUC)

For Figure 1, conditioned media was harvested from hiPSC-CMs cultured in T175 flasks and 6-well plates. For Figure 4, 2-3 million hiPSC-CMs were grown per well in 6-well plates and 3 full plates per condition were stimulated for 48 hours with 2 ml per well of maturation media only (control), 10 ng/ml TNF-α or exposed to hypoxia by placing the plate in a BD GasPak™ EZ Anaerobe Pouch System (BD Biosciences, 260683). After 48 hours, the conditioned medium was collected and sEV isolation was performed using dUC. To remove cell debris, the conditioned medium was centrifuged for 5 minutes at 300 xg and for 10 minutes at 2000 xg. Next, the supernatant was added to ultracentrifuge tubes (Beckman Coulter, 344058) and centrifuged for 30 minutes at 10 000 xg followed by centrifugation for 70 minutes at 100 000 xg. The pellet of the last step was resuspended in 50 µl of PBS. All centrifugation steps were done at 4°C and the last two steps were performed using Optima XE-90 ultracentrifuge (Beckman Coulter, A94471) with rotor SW32Ti (Beckman Coulter, 369694).

### Size exclusion chromatography (SEC)

Conditioned medium of hiPSC-CMs expressing HA-NL-tetraspanin reporters was harvested and centrifuged for 5 minutes at 500 xg to remove cell debris. Separation of sEVs from soluble proteins was performed using qEVoriginal 70 nm SEC column (IZON, ICO-70) according to the manufacturer’s instructions. 1.5 ml of supernatant was added on top of the SEC column and allowed to enter by gravity. PBS was used as running buffer and fractions of 0.5 ml were collected manually up to 26 fractions. 50 µl of each fraction was added in duplicate into a white 96-well plate (Greiner Bio-One, 655075) and 50 µl of Nano-Glo reagent (Promega, N1110, 1:1000) was added in each well using a multichannel. Luminescence activity was measured after 3-minute incubation using SpectraMax iD3 microplate reader (Molecular Devices) and analyzed using SoftMax Pro 7 Software (Molecular Devices).

### Protein quantification and western blot

EV samples were prepared by lysing EVs directly in concentrated RIPA buffer (Millipore, 20-188) and protease/phosphatase inhibitors (Cell Signaling Technology, 5872). Cells were collected in 1x RIPA buffer containing protease/phosphatase inhibitors and incubated on ice for 30 minutes. Next, cell lysate was centrifugated at 12 000 xg for 15 minutes at 4°C and the resultant supernatant was collected and stored at −80°C. Protein concentration was determined using the micro-BCA protein assay kit (Thermo Scientific, 23235) according to the manufacturer’s instructions. The protein concentration of cell lysates was measured by diluting samples 1:50 in PBS. The protein content of SEC fractions was quantified by adding 50 µl of each SEC fraction in duplicate into a 96-well plate except for protein-enriched fractions (usually starting on fraction 16), which were diluted 1:25. Measurements were done using Multiskan FC microplate photometer (Thermo Scientific, 51119000) and analyzed with SkanIt software version 3.1 (Thermo Scientific, 5187139). To analyze proteins with western blot, EV samples from equal numbers of cells and equal volumes of conditioned medium were loaded without further dilution. 5 µg of cell lysate was used. Samples were prepared in NuPAGE LDS loading buffer (Invitrogen, NP0007) and reduced using NuPAGE reducing agent (Invitrogen, NP0009), followed by boiling at 95°C for 10 minutes. For CD63 detection, non-reduced samples were used. Subsequently, samples were loaded into Bolt 4-12% Bis-Tris Plus 1.0mm x 15 well gels (Invitrogen, NW04125BOX), and PageRuler Plus (Invitrogen, 26619) was used as a protein ladder. Electrophoresis was done at 100 volts for 75 minutes in NuPAGE MOPS SDS running buffer (Invitrogen, NP0001). Gels were transferred to a polyvinylidene difluoride (PVDF) membrane (Invitrogen, IB24001) using the iBlot2 system for 7 minutes at 20 volts. Membranes were blocked in TBS containing 5% BSA (Roche, 10735086001) for 1 hour at room temperature or overnight at 4°C. Primary antibody incubation was done overnight at 4°C in TBS-0.1%Tween20 (Sigma-Aldrich, 822184) containing 0.5% BSA. Primary antibodies were used against CD9 (Merck Millipore, CBL162, 1:1000), CD63 (BD Biosciences, 556019, 1:1000), CD81 (Santa Cruz, sc-166029, 1:500-1000), syntenin (OriGene, TA504796, 1:1000), annexin A1 (Abcam, ab214486, 1:1000),

GAPDH (Cell Signaling Technology, 2118, 1:1000), GM130 (BD Biosciences, 610823, 1:500), calnexin (GeneTex, GTX101676, 1:1000) and HSP70 (Biolegend, 648001, 1:500). Secondary antibodies were diluted 1:7500 in a mixture of 50% v/v of Intercept blocking buffer (LI-COR, 927-60001) in TBS-0.1%Tween20 and incubated for 2 hours at room temperature in the dark. Secondary antibodies included Alexa 680-conjugated goat anti-mouse (Invitrogen, A21057) and IRDye 800CW-conjugated goat anti-rabbit (LI-COR Biosciences, 926-32211). Imaging was achieved using channels 700 nm and 800 nm of the Odyssey M (LI-COR Biosciences) and processed with LI-COR acquisition software (LI-COR Biosciences). Washing steps were performed between antibody incubations 3 times for 10 minutes in TBS-0.1%Tween20 buffer.

### Nanoparticle tracking analysis (NTA)

NTA was conducted using NanoSight NS500 (Malvern Panalytical). To study particle concentration and size distribution of isolated sEVs, EV samples were diluted in PBS to obtain a particle concentration of 30 to 100 particles per frame. Particle concentration of pooled SEC fractions was measured without dilution, except for protein-enriched fractions (21-26) that were diluted 1:5 in PBS. Five videos of 60 seconds, or 30 seconds in the case of pooled SEC fractions, were captured using a camera level of 16 and a detection threshold of 6. Videos were analyzed using NanoSight NTA 3.3 software (Malvern Panalytical).

### Transmitted electron microscopy (TEM)

To image sEVs isolated by dUC, the EV sample was added to a carbon-coated grid (75–200 mesh) for 15 minutes. Unbound vesicles were removed with PBS washing followed by fixation with 1% glutaraldehyde in PBS buffer for 30 minutes. Next, samples were counterstained with 2% uranyl-oxalate for 10 minutes at room temperature and embedded in a mixture of 1.8% methylcellulose and 0.4% uranyl acetate for at least 10 minutes at 4°C. Grids were dried for 1 hour at room temperature. sEVs were imaged on JEM 1011 TEM microscope (Jeol), equipped with a tungsten filament and a DCC CMOS camera Phurona (Emsis) with a resolution of 12 megapixels. The EM beam was set to a voltage of 80 KV.

### Plasmids

The pLenti6.3-CD63-pHluorin and pLenti6.3-HA-NL-CD63 lentiviral plasmids were described previously.^25,68^ To generate HA-NL-CD9 and HA-NL-CD81 lentiviral plasmids, CD9 and CD81 cDNA sequences were amplified using the primer pairs 5’ - ataggatccatgccggtcaaaggaggcacc – 3’ + 5’ – atatctagagctagaccatctcgcggttcc – 3’, and 5’ – ataggatccatgggagtggagggctgcacc – 3’ + 5’ – atatctagagctagtacacggagctgttcc – 3’, respectively, to add BamHI and XbaI restriction sites. The amplified CD9 and CD81 fragments were cloned into BamHI/XbaI-digested HA-NL-CD63-pCMV-Sport6 vector, to generate HA-NL-CD9 and HA-NL-CD81, which were subsequently subcloned into the pLenti6.3/TO/V5-DEST vector (Thermo Fisher Scientific).

### Generation of HA-NL-tetraspanin and CD63-pHluorin expressing hiPSC-CMs

HA-NL-tetraspanin and CD63-pHluorin expressing hiPSC-CMs were produced using lentiviral transduction. Lentivirus production was done in HEK293FT cells, which were seeded in 10 cm dishes coated with 0.1% gelatin (Sigma-Aldrich, G1890) and cultured in DMEM (Gibco, 41965039) supplemented with 10% FBS (Corning, 35-079-CV) and 1% penicillin-streptomycin (Gibco, 15140122). HEK293FTs were transfected with plasmid DNA using lipofectamine 3000 (Invitrogen, L3000015) at 70-80% confluency. 3.7 μg pLenti6.3-HA-NL-tetraspanin plasmids or pLenti6.3-CD63-pHluorin plasmid were co-transfected with 0.9 μg pREV (Addgene, 12253) and 1.8 μg pRRE (Addgene, 12251) packaging plasmids, and 1.1 μg pMD2.G (Addgene, 12259) lentiviral envelope plasmid. P3000 reagent was added to the DNA mix in a 2 μl to 1 μg ratio. Next, plasmids were complexed with lipofectamine 3000 using a 1 μg to 1.5 μl ratio in Opti-MEM medium (Gibco, 31985070) for 15 minutes at room temperature.

Subsequently, the complexes were added on top of HEK293FT cells after removing half of the culture medium and incubated for 18 hours before changing the media. Lentivirus-containing media was collected 48-72 hours after transfection. Cell debris was removed by centrifugation at 500 xg for 5 minutes and filtration using a 0.45 µm SCFA syringe filter (Corning, 516-1954), followed by aliquoting and storage at −80°C until further use. To produce stable HA-NL-tetraspanin expressing hiPSC-CMs, cells were cultured in T175 flasks and 7 ml of lentivirus-containing media was added on top of 20 ml of expansion media RPMI supplemented with B27 (Gibco, 17504044) and 2 µM CHIR99021 (SelleckChem, S2924), when cells reached 60-70% confluency. After 48 hours, the antibiotic selection was performed for at least 5 days using expansion media containing 5 µg/ml blasticidin S (Gibco, A1113903). hiPSC-CMs were kept in expansion media for another 2-3 days to allow cell recovery and then cultured in maturation media for at least 21 days before split into matrigel-coated white 96-well plates (Greiner Bio-One, 655083) to perform NanoLuc assay. To generate stable CD63-pHluorin expressing hiPSC-CMs, cells were grown in a 6-well plate and cultured for 5 days in expansion medium containing 2 ml of lentivirus-containing medium per well. Antibiotic selection was done in 5 µg/ml blasticidin for 4 days and cells were kept in expansion media for 3 days before starting the maturation process. After 21 days, hiPSC-CMs were passed into matrigel-coated CELLview slides (Greiner Bio-One, 543078) to allow live TIRF imaging.

### Immunofluorescence microscopy

hiPSC-CMs were grown in CELLview slides (Greiner Bio-One, 543078) and fixed using 4% PFA for 15 minutes, followed by washing with PBS. Cells were permeabilized with 0.1% Triton X-100 (Sigma-Aldrich, T9284) for 5 minutes, washed with PBS, and blocked in 5% BSA (Roche, 10735086001) in PBS solution. Subsequently, primary antibody incubation was performed in blocking buffer with anti-HA-tag (rat, Roche, 11867423001, 1:250) or anti-CD9 (mouse, Merck Millipore, CBL162, 1:100) or anti-CD63 (mouse, BD Biosciences, 556019, 1:100) and anti-cardiac troponin (rabbit, Abcam, ab45932, 1:300). For Figure S1D, hiPSC-CMs were grown on coverslips and were stained with anti-α-actinin (mouse, Sigma-Aldrich, A7811, 1:300) and anti-Tomm20 (rabbit, Santa Cruz, sc-11415, 1:300). Both blocking and primary antibody incubations were done for 1 hour at room temperature or overnight at 4°C. Secondary antibody incubation was done together with the nuclei staining using Hoechst 33342 (Invitrogen, H1399, 1:10 000) for 1 hour at room temperature in the dark. Alexa Fluor 647-linked anti-rat (goat, Invitrogen, A21247) and Alexa Fluor 488-linked anti-rabbit (donkey, Invitrogen, A21206), or Alexa Fluor 488-linked anti-mouse (donkey, Invitrogen, A21202) and Alexa Fluor 647-linked anti-rabbit (donkey, Invitrogen, A31573), or Alexa Fluor 488-linked anti-mouse (goat, Invitrogen, A11029) and Alexa Fluor 568-linked anti-rabbit (goat, Invitrogen, A11011), were used as secondary antibodies and were diluted 1:400 in blocking buffer. Washing steps after each antibody incubation were done 3 times for 10 minutes with PBS. Cells were kept in PBS until imaging with the LSM700 confocal microscope (Zeiss) using ZEN 2011 software (Zeiss). Image acquisition was performed using an oil 63x/1.40 objective and the 405, 488, and 633 nm laser lines. Images were analyzed using Fiji software version 2.14.^102^ For Figure S1D, samples were imaged with 20x and 40x objectives of EVOS™ M7000 Imaging System (Thermo Fisher Scientific).

### HA-NL-tetraspanin secretion assay

HA-NL-tetraspanin hiPSC-CMs were seeded in a white 96-well plate (Greiner Bio-One, 655083) at a density of 70 000 cells per well. Drug incubations were performed in 200 µl of maturation media per well and in the presence of 10 ng/ml TNF-α when mentioned. Nutrient starvation was carried out by culturing the cells in DMEM without glucose (Gibco, 11966025) and without supplements. To expose the cells to hypoxia, plates were incubated in a BD GasPak™ EZ Anaerobe Pouch System (BD Biosciences, 260683), placing one plate per bag. After the specified time point, 100 µl of conditioned medium was collected into a transparent U-bottom 96-well plate using a multichannel and centrifuged at 500 xg to remove any floating cells from the supernatant. Next, 50 µl of supernatant was transferred into a white 96-well plate (Greiner Bio-One, 655075) and 50 µl of Nano-Glo reagent was added using a multichannel and incubated for 3 minutes. Nano-Glo assay reagent consisted of a 1:1000 dilution of Nano-Glo substrate (Promega; N1110) in buffer (100mM 2-(N-Morpholino)ethanesulfonic acid hydrate (M8250; Sigma-Aldrich), 1mM trans-1,2-Diaminocyclohexane-N,N,Nʹ,Nʹ-tetraacetic acid monohydrate, 150mM KCl, 1mM DL-Dithiothreitol (D0632; Sigma-Aldrich), 35mM Thiourea (T7875; Sigma-Aldrich) and 0.5% Tergitol NP-40 (NP40S; Sigma-Aldrich), pH 6.). Luminescence activity was measured using CLARIOstar Plus (BMG Labtech) and analyzed with MARS Data Analysis Software (BMG Labtech). The same was performed on the original plate containing the donor cells by adding 100 µl of Nano-Glo reagent per well. HA-NL-tetraspanin secretion was calculated by first dividing the luminescence of the supernatant with that of the cells, followed by normalization to the DMSO control of each plate.

### Live TIRF microscopy

TIRF imaging was performed as previously described.^68^ In short, hiPSC-CMs transduced with CD63-pHluorin were seeded in CELLview slides (Greiner Bio-One, 543078) and exposed to maturation media only (control), 10 ng/ml TNF-α or incubated in a BD GasPak™ EZ Anaerobe Pouch System (BD Biosciences, 260683) to mimic hypoxic condition. After 48 hours, hiPSC-CMs were imaged by live TIRF microscopy. A Nikon Eclipse Ti with Perfect Focus System setup was used with a laser bench coupled into a Nikon Apo TIRF 100x N.A. 1.49 oil objective. Images were acquired at 2 Hz, using a Photometrics CoolSNAP HQ2 CCD. Live-imaging experiments were performed in maturation medium at 37°C and 5% CO2 using a Tokai Hit INUBG2E-ZILCS Stage Top Incubator. Images were acquired using MetaMorph software (version 7.10.2.240) and fusion activity was quantified as described before.^68^

### Cell viability assay (alamarBlue)

Cell viability was performed using alamarBlue reagent (Invitrogen, DAL1025) diluted 1:10 in maturation media followed by filtration with a 0,22 µm CME syringe filter (Carl ROTH, KH54.1). Incubation with small molecule inhibitors, AngII and TNF-α was done in a transparent 96-well plate for 48 hours using the same concentrations as used for the HA-NL-tetraspanin secretion assays. After 48 hours, the medium was replaced by 100 µl alamarBlue solution per well and incubated overnight. alamarBlue solution was also added in 6 empty wells as a blank to control for fluorescence background. The next day, 50 µl of conditioned medium was transferred to a black 96-well plate. Fluorescence was measured at an excitation wavelength of 560 nm and an emission of 590 nm in CLARIOstar Plus (BMG Labtech) and analyzed using MARS Data Analysis Software (BMG Labtech). Cell viability was calculated by subtracting the fluorescence background of the blank (no cell incubation) from all conditions and normalizing to the DMSO control.

### Statistical analysis

Data was analyzed and visualized using Prism 10.0 software (GraphPad). The sample number and statistical tests used are indicated in the figure legends. Statistical significance is denoted by asterisks with *p < 0.05, **p < 0.01, ***p < 0.001, ****p < 0.0001

## Supplementary information

**Figure S1.**
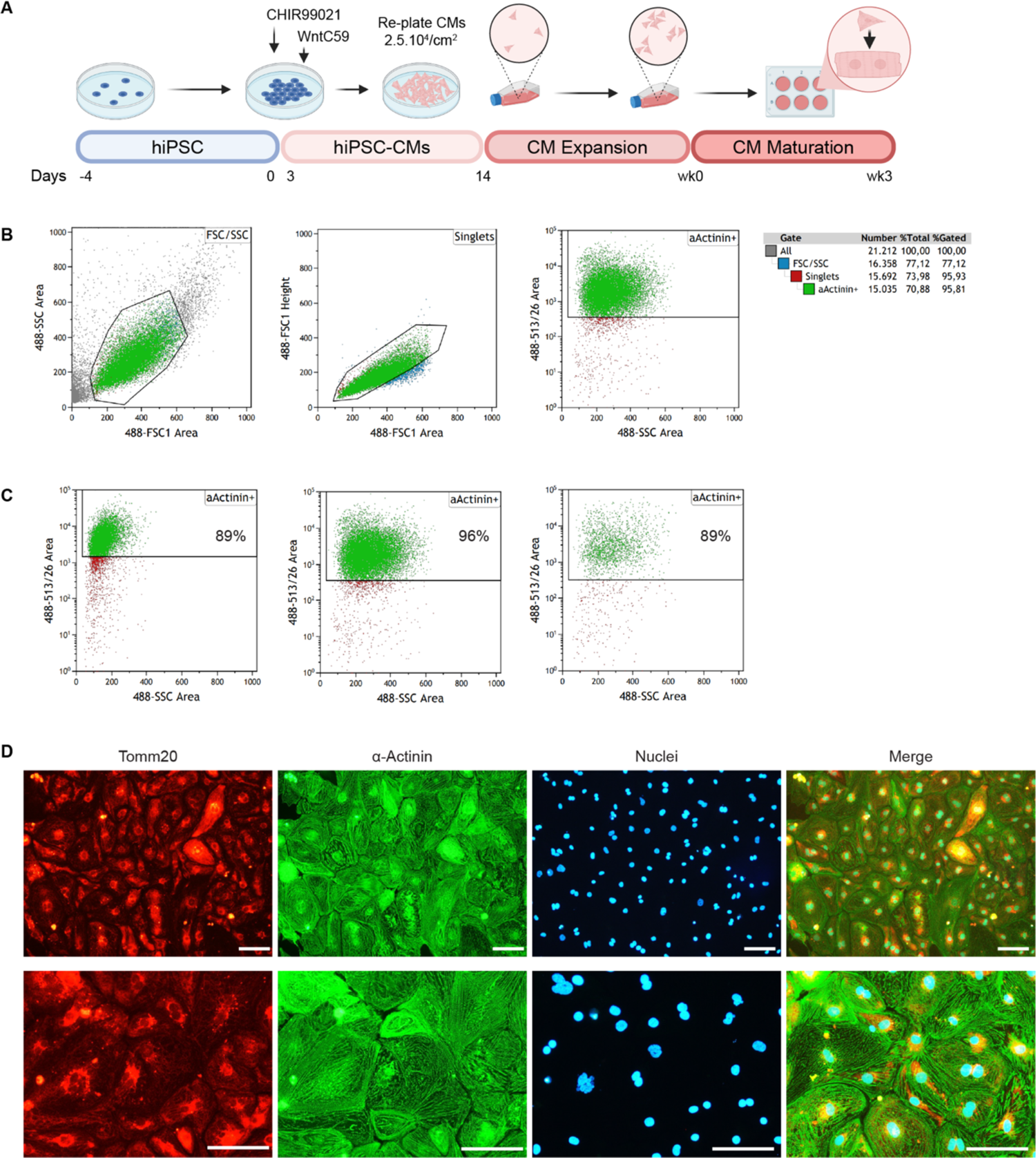
Generation and purity control of hiPSC-CMs. **A**. Schematic workflow of the hiPSC-CM generation and culturing. **B**. Representative gating strategy to select alpha-actinin positive (α-actinin+) hiPSC-CMs. Abbreviations: SSC, side scatter; FSC, forward scatter. **C**. Flow cytometry analysis of α-actinin+ hiPSC-CMs of three individual differentiations as previously described.^47^ **D**. Representative immunofluorescence images of hiPSC-CMs after expansion and stained for Tomm20 (mitochondria) and α-actinin (cardiac sarcomeres). Scale bar = 100 µm.

**Figure S2.**
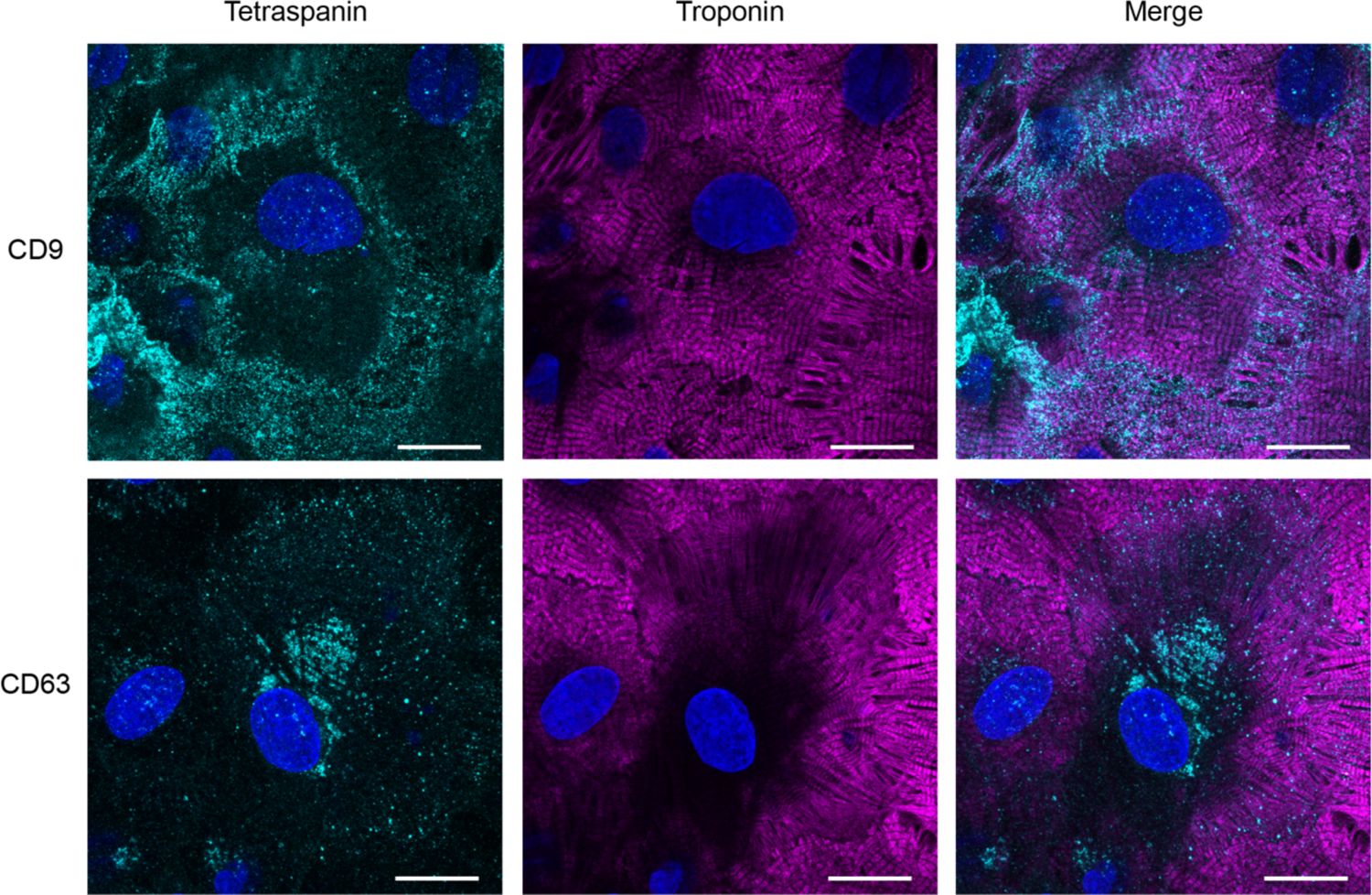
Representative immunofluorescence images of hiPSC-CMs after maturation and stained for CD9 or CD63 and cardiac troponin. Scale bar = 20 µm.

**Figure S3.**
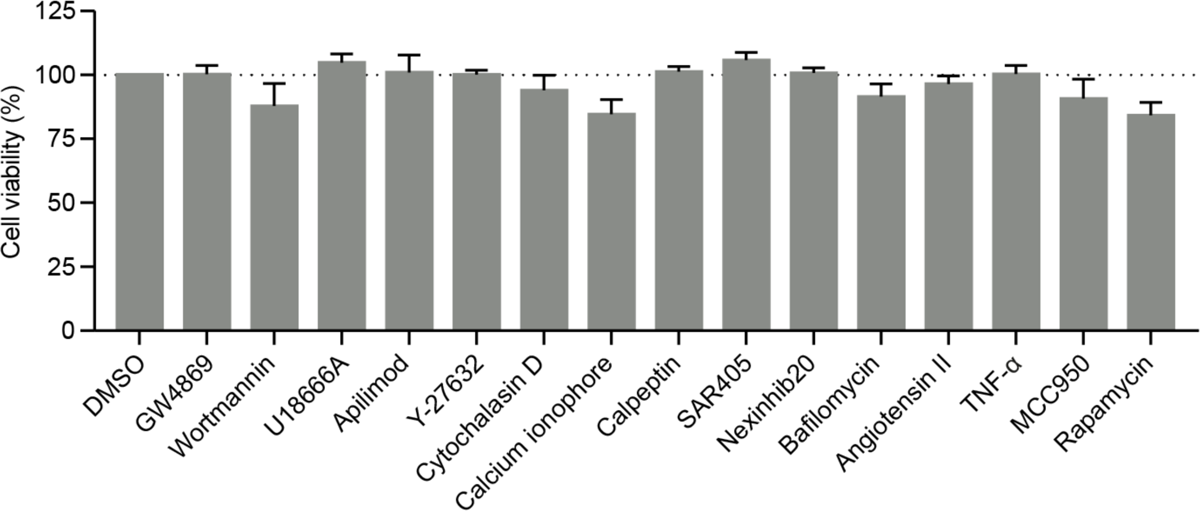
Cell viability (alamarBlue assay) of hiPSC-CMs after exposure to small molecules or proteins used throughout the study. Results are plotted as mean ± SEM (n = 3).

**Figure S4.**
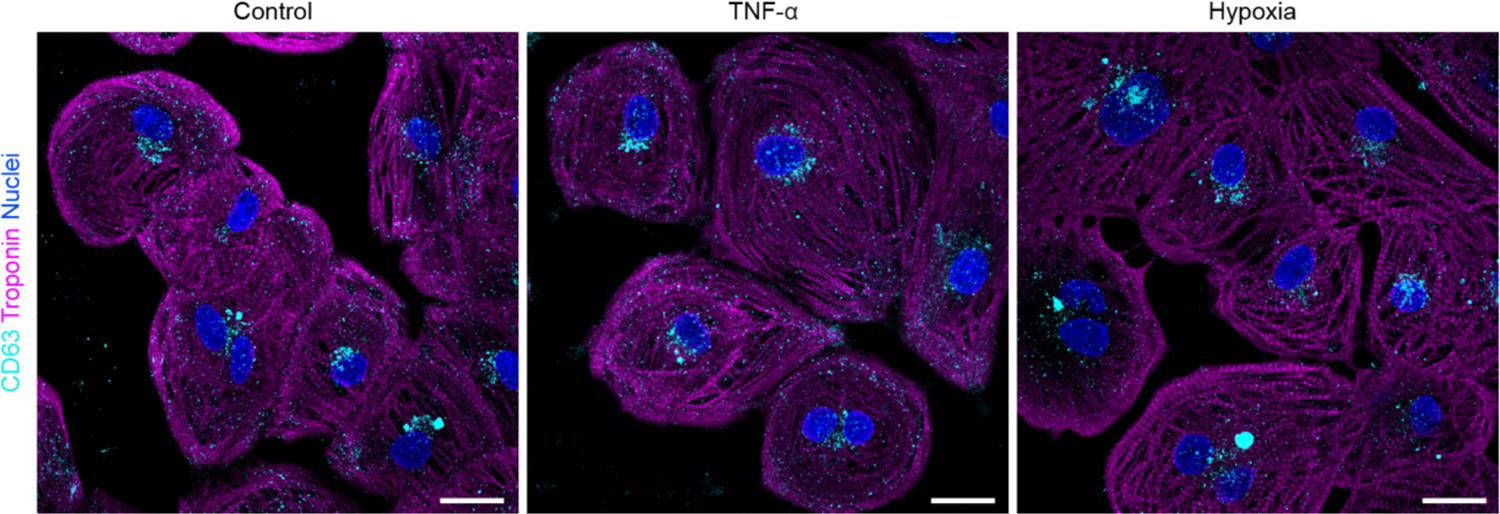
Representative immunofluorescence images of hiPSC-CMs upon incubation for 48 hours of control, TNF-α, and hypoxic conditions and stained for CD63 and cardiac troponin. Scale bar = 20 µm.

---

**Video S1.** Morphology of beating hiPSC-CMs 26 days after maturation imaged using a 10x objective.

**Video S2.** TIRF microscopy of hiPSC-CMs expressing CD63-pHluorin. The video is played at 10x speed. Scale bar 10 μm.

**Video S3.** TIRF microscopy of hiPSC-CMs expressing CD63-pHluorin after 48 hours incubation with TNF-α. The video is played at 10x speed. Scale bar 10 μm.

**Table S1.** Small molecules used during this study and their effect on extracellular vesicle (EV) biogenesis pathways. Abbreviations: vATPase, vacuolar ATPase; nSMase2, neutral sphingomyelinase 2; PI3K, phosphatidylinositol 3-kinase; NPC-1, Niemann-Pick disease type C1; PIKfyve, FYVE finger-containing phosphoinositide kinase; ROCK, Rho-associated protein kinase; JFC1, Synaptotagmin like 1; ILV, intraluminal vesicles; MVBs, multivesicular bodies.

| Name | Target | Effect on EV secretion | Ref. |
| --- | --- | --- | --- |
| Bafilomycin | vATPase | Inhibits vATPase-mediated late endosomal/lysosomal acidification, stimulating exosome secretion. | 54 |
| GW4869 | nSmase2 | Inhibits nSmase2-mediated ILV formation, decreasing exosome release. Reported to stimulate ectosome secretion. | 64,92 |
| Wortmannin | PI3K | Inhibits PI3K-mediated ILV formation, preventing MVB formation and exosome release. | 103 |
| U18666A | NPC-1 | Inhibits cholesterol transport from the lysosome, decreasing exosome secretion. | 80,104 |
| Apilimod | PIKfyve | Impairs fusion of lysosomes with MVBs and autophagosomes, inducing secretory autophagy and increasing exosome secretion. | 86 |
| Y-27632 | ROCK | Inhibits ROCK-mediated outward budding of the plasma membrane via actomyosin remodeling, decreasing ectosome release. | 77,78 |
| Cytochalasin D | Inhibitor of actin polymerization | Increases EV release potentially by reducing restrictions on exocytosis. | 26 |
| Calcium ionophore A23187 | Calcium | Increases intracellular calcium concentration, increasing EV secretion. | 105 |
| Calpeptin | Calpain 1 and 2 | Inhibits Calpain-mediated cytoskeleton remodeling, preventing ectosome release. | 87,88 |
| SAR405 | VPS34 | Inhibits VPS34-mediated canonical autophagy, reducing potentially amphisome-dependent exosome secretion. | 106 |
| Nexinhib20 | Inhibitor of the Rab27a-JFC1 interaction | Inhibits Rab27a-mediated docking of MVBs, decreasing exosome secretion. | 79 |

## Notes

### Competing Interest Statement

The authors have declared no competing interest.

